# Thermal fertility limits reveal underestimated climatic suitability for *Aedes aegypti* in temperate regions

**DOI:** 10.64898/2026.08.06.743224

**Authors:** A Haziqah-Rashid, Soeren Metelmann, Kinga Stobierska, Katherine Gawne, Huilan Yu, Ken Sherlock, Matthew Baylis, Ewa Chrostek, Marcus Blagrove

## Abstract

Predicting species distributions under climate change typically relies on thermal limits for survival. However, recent evidence suggests that sublethal temperatures that reduce fertility can constrain distributions more strongly than temperatures causing mortality alone. Despite the importance of mosquitoes as vectors of human disease, thermal fertility limits remain largely uncharacterized in these insects. Here, we show that fertility in *Aedes aegypti* is highly sensitive to thermal stress and exhibits distinct sex-specific responses. Adult males were more vulnerable to both heat and cold exposure than females. At 38°C, survival remained high in both sexes despite significant reproductive impairment, indicating that sterility can occur independently of mortality. Similarly, exposure to 2°C induced male sterility whereas females maintained reproductive function. Morphological analyses of reproductive organs showed temperature-associated damage consistent with the observed fertility loss. Incorporating these fertility thresholds into ecological niche models generated different predictions of climatic suitability compared with models based solely on lethal temperature limits. Our findings demonstrate that sublethal effects on reproduction can substantially influence estimates of mosquito climatic suitability and highlight the importance of incorporating fertility based thermal limits into projections of vector distributions and disease risk under future climate change.

## INTRODUCTION

The yellow fever mosquito, *Aedes aegypti*, is native to sub-Saharan Africa and is now widely distributed across tropical and subtropical regions, with established populations extending into some warm temperate areas (Powell and Tabachnick, 2013). This species is a primary vector for dengue virus, which continues to pose a significant public health threat worldwide. The number of reported cases of dengue increased dramatically to about 15 million in 2024, compared with only half a million cases in 2000. Additionally, from January to July 2025, more than 4 million cases and over 3,000 deaths were reported across 97 countries in Africa, the Americas, Europe, Southeast Asia, and the Western Pacific (World Health Organization, 2025).

Mosquito development and survival are heavily influenced by environmental factors. For example, *Ae. aegypti* eggs can endure prolonged dry conditions by entering a dormant state and adjusting protein expression to survive desiccation (Prasad et al., 2023). Temperature plays a crucial role, alongside other factors such as humidity, rainfall, and elevation, in shaping mosquito behaviour and life history traits (Field et al., 2022).

As global temperatures rise, the geographical distribution of mosquito species is shifting, with both empirical observations and predictive models indicating continued expansion into newly suitable regions. Recent studies predict a poleward expansion of *Ae. aegypti* and *Ae. albopictus* (Laporta et al., 2023) and modelling suggests this will continue in response to climate change, with mosquitoes gradually colonising new areas as conditions become thermally suitable. These changes have already been observed. Following the disappearance of from much of mainland Europe and the Mediterranean region during the mid-twentieth century, the species was reintroduced to Madeira, Portugal in 2005 where it has since established a persistent population (Seixas et al., 2019; Schaffnerr and Matthis, 2014). Numerous predictive models have attempted to map the future spread of these vectors, incorporating temperature thresholds, life cycle dynamics, land use, and precipitation patterns (Iwamura et al., 2020; Kraemer et al., 2015; Kamal et al., 2018). For example, Iwamura et al. (2020) predicted that the global climatic suitability for *Ae. aegypti* life-cycle completions (i.e. complete egg-to-adult development cycles) would increase by approximately 3.2-4.4% per decade by 2050, compared with an observed increase of approximately 1.5% per decade between 1950 and 2020. It is also predicted that overall climatic suitability for *Ae. aegypti* in Europe will remain relatively low, but southern regions particularly across the Iberian Peninsula, Italy and Greece are projected to support high levels of life cycle completion (Iwamura et al., 2020). While temperate regions in the Northern Hemisphere are becoming more suitable for *Aedes* mosquitoes, some tropical regions, particularly in the Southern Hemisphere, may see a reduction in range as temperatures exceed optimal thresholds for mosquito persistence and precipitation declines (Ryan et al., 2019; Laporta et al., 2023).

Despite growing interest in forecasting mosquito distributions, most predictive models rely on survival limits rather than sterility thresholds. Experimental studies in *Drosophila* have demonstrated that thermal fertility limits (TFL) may provide a more sensitive predictor of thermal tolerance than survival-based limits alone (Walsh et al., 2019; Parratt et al.,2021). However, TFL remain largely uncharacterised in mosquitoes and other disease vectors, limiting our understanding of how reproductive constraints shape mosquito distributions under changing climates. Furthermore, previous studies have largely focused on a single sex, despite evidence that thermal stress can differentially affect male and female fertility (Walsh et al., 2019). Investigating thermal fertility responses in both sexes is important for accurately characterising mosquito thermal limits and predicting population persistence.

This study explored how temperature affects *Ae. aegypti* beyond survival alone. Specifically, we investigated critical thermal limits (CTL) and thermal fertility limits in both male and female mosquitoes together with the mechanisms underlying thermally induced sterility. Finally, these experimentally derived thermal limits were integrated into updated distribution models and risk maps to improve predictions of future *Aedes* spread under currently accepted climate change scenarios.

## METHODOLOGY

### Mosquito rearing

Liverpool strain *Aedes aegypti* eggs were hatched overnight in 200 mL of dechlorinated tap water containing yeast prepared by dissolving one yeast tablet in 1 L of water, at 27°C. First-instar larvae were transferred to rearing trays containing dechlorinated tap water that had been left to stand overnight, at a standardized density of 200 larvae per 2 L. Trays were maintained at 27°C and 80-85% relative humidity, with larvae fed TetraMin fish tablets on alternate days, and water replaced as required. Pupae were transferred to emergence containers and placed in cages under a 13:11 h light: dark photoperiod with 10% sucrose provided *ad libitum*. Adults were sugar-starved for 24 h prior to weekly blood feeding using human blood and a Hemotek feeder. Oviposition substrates – filter papers - were provided, and eggs were collected after 5 days and dried at 27°C and 80% relative humidity for one week before hatching.

### Pupae sexing for experiments

To ensure adults were unmated, mosquitoes were sexed at the pupal stage and sex confirmed after emergence using standard morphological characteristics. Individuals were placed in cages separately to prevent mating. Damaged pupae and adults were excluded from experiments.

### Heat and cold shocking experiments

#### a) Mortality experiments

15 male and 15 female adults, aged 3 – 5 days, were exposed to high (27, 38, 41, 43, 46, 47°C) and cold (-10, -2, -1, 0, 2, 4, 6, 8°C) temperatures for 4 and 6 hrs respectively. The exposure duration at high temperatures was determined based on the protocol by Bretman *et al*. (2024) while cold exposure duration was based on preliminary trials. No sugar solution was provided during the exposure period. After exposure, adult mosquitoes were allowed to recover at 27°C overnight. Mortality was assessed 24 h post-exposure. Each temperature treatment was replicated four times.

#### b) Sterility experiments

Surviving adults were paired with non-thermally shocked, unmated counterparts at a ratio of one treated individual to three untreated individuals of the opposite sex. For male fertility assays, one treated male was placed with three untreated virgin females, whereas for female fertility assays, one treated female was placed with three untreated virgin males. Adults were allowed to mate for four days and then blood-fed using the same Hemotek feeding protocol for colony maintenance. Filter papers were then provided, and eggs were collected after five days. Eggs were dried for five days before hatching using the same protocol describe for colony maintenance. Fertility was assessed from the total number of eggs laid per cage and the corresponding egg hatch rate.

### Survival and sterility calculations

Survival rates were assessed 24 h post-exposure. Hatching success is commonly calculated as the number of larvae hatched divided by the total number of eggs laid (Supplementary Fig. 1.1). However, when few adults survive but produce large egg batches, this approach can overestimate fertility relative to the number of adults exposed. To address this limitation, fertility was additionally calculated using a binary metric based on the presence or absence of larval hatch per exposed adult. Fertility percentage was calculated as:

**(Number of adults with hatched larvae / Total number of adults exposed) x 100**

We present both sets of results (Supplementary Figure 1.1), showing that neither calculation method is entirely perfect.

### Morphology observations

To assess whether thermal exposure caused morphological changes to the reproductive organs, all surviving mosquitoes from each temperature treatment and the control (27°C) were dissected. Dissections were performed in phosphate-buffered saline (PBS) under a stereomicroscope (Motic, SMZ-171-Binocular LED) and the reproductive organs were examined using a phase contrast microscope (Olympus, CX21) at 100× total magnification (10× objective x 10× eyepiece). The organs examined in females were the ovaries and spermathecae, while accessory glands were observed in males. Morphological changes were assessed qualitatively by visual comparison between the temperature-treated and control mosquitoes. Control adults were classified as either mated and fertile or unmated and sterile for comparison. Representative images presented in the results were selected from individuals classified as fertile, partially sterile (less than 50% egg hatch), or sterile based on their corresponding egg hatch rate. In addition, male accessory glands were also examined because they play an important role in stimulating ovulation in female *Ae. aegypti*.

### Data analysis

Data analyses were run using the statistical software R, version 4.2 (R Development Core Team, 2022), except where stated. Thermal performance curves for survival and fertility were fitted using nonlinear least-squares regression (nlsLM function, minpack.lm package) with a Gaussian-type model (formula can be found in the Supplementary Method). The area between survival and fertility curves was shaded to represent the thermal fertility gap.

Probit analyses were done using SPSS to calculate the regression for lethal temperature (CTL_5_ – CTL_95_) and the sterility temperature (TFL_5_ – TFL_95_). 5 and 95 represent the temperature that is causing 5% and 95% of the population to be dead or sterile. These data were then incorporated into the model for the lowest and highest temperatures that can cause lethality and sterility.

### Risk map modelling

Both mortality and sterility data from all replicates were subjected to Probit analysis. Critical thermal limit (CTL) which resulted in 5% - 95% (CTL_5_ – CTL_95_) values were calculated from a log dosage-probit mortality regression line using SPSS (Version 27). We then mapped regions that are too warm or too cold for *Ae. aegypti* to live and to reproduce: *Ae. aegypti* occurrence data (Kraemer et al., 2015) were combined with geographic, socioeconomic and climatic data (Gridded Population of the World (GPWv4), WorldClim and NASA Earthdata) to determine the ecological niche for *Ae. aegypti*. The results of the probit analysis are used to restrain the ecological niche further. The maximum temperature of the warmest month (bioclim 05) was used to indicate regions that are too warm for survival or reproduction with the cutoffs CTL90 and TFL90 (Parratt et al., 2021). Similarly, the minimum temperature of the coldest month (bioclim 06) was used to indicate regions that are too cold for survival or reproduction. These steps were repeated with climate projections for the 2050s to indicate how suitable areas might expand or shift with a changing climate.

## RESULTS

### Effects of high and low temperatures on adult *Ae. aegypti*

We found a significant gap between the lethal and the sterility temperature in both male and female adults (Figures 1 and 2), particularly close to thermal extremes. In general, the number of adult survivors increased as the temperature increased from -2°C (following a 6 hrs cold exposure) and declined at high temperature (38°C) and reached complete lethality at 47°C. Sterility also increased at both high and low temperatures, but its relationship with survival differed between cold and heat stress. At low temperatures (0 - 8°C), a high proportion of adults survived the prolonged exposure but remained largely sterile and produced no viable offspring. In contrast, fertility was highest at 27 – 35°C and declined sharply at 38°C, where more than half of the surviving population was sterile despite approximately 75% adult survival. Sex-specific responses were also evident where males (Figure 2) showed higher mortality and sterility at cold temperatures compared to females (Figure 1), whereas at high temperatures males were more likely to die while females survived but were sterile.

**Figure 1:**
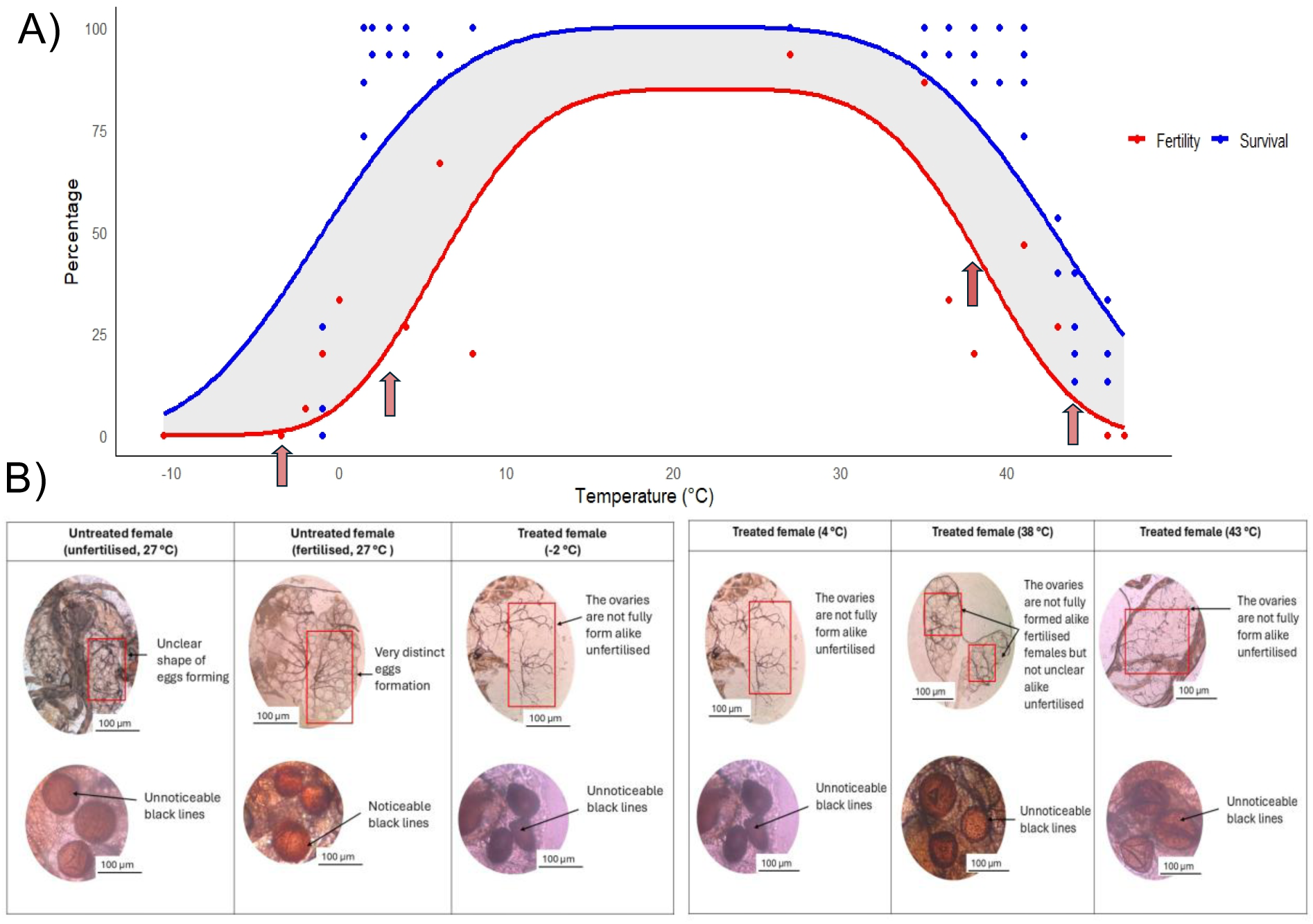
*Ae. aegypti* females exposed to high and low temperatures: Dots represent four replicates for each thermal treatment. One replicate consisted of 15 mosquitoes. Red and blue lines represent fertility and survival, respectively. Grey shaded areas between the lines show the thermal fertility gap. A) Adult females exposed to temperatures ranging from -10 to 47°C. B) Morphology of untreated fertilised and unfertilised females maintained at 27°C (control) and females exposed to −2, 4, 38, and 43°C, respectively, as indicated by the red arrows on the graph, corresponding to temperatures near the lower and upper thermal fertility limits. Temperatures selected for ovary imaging represent conditions associated with reduced fertility at the lower and upper ends of the fertility curve. Images are shown at 100× magnification. The top images show ovaries and the bottom images show spermathecae. Black lines visible in the spermathecae indicate the circular orientation of the spermatozoa which can be observed clearly in fertilised females but are absent in unfertilised and treated females. Red boxes highlight regions of ovarian development and egg formation. Scale bar = 100 µm.

**Figure 2:**
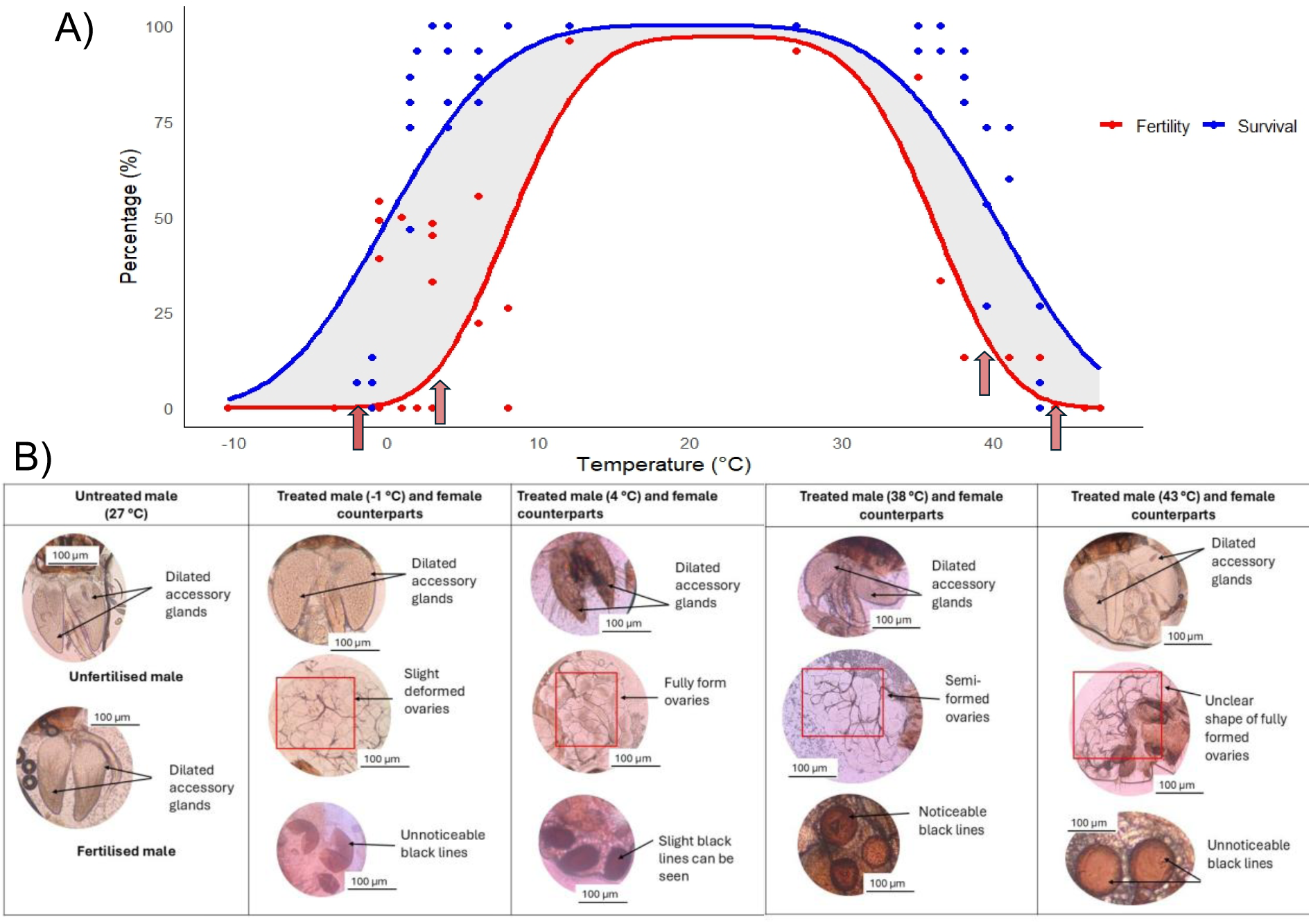
*Ae. aegypti* males exposed to high and low temperatures: Dots represent four replicates for each thermal treatment. One replicate consisted of 15 mosquitoes. Red and blue lines represent fertility and survival, respectively. Grey shaded areas between the lines show the thermal fertility gap. A) Adult males exposed to temperatures ranging from -10 to 47°C. B) Morphology of untreated males maintained at 27°C (control) and males exposed to −1, 4, 38, and 43°C and their female counterparts, respectively, as indicated by the red arrows on the graph. Temperatures selected for imaging represent conditions associated with reduced fertility at the lower and upper ends of the fertility curve. Images are shown at 100× magnification. The upper images show male accessory glands, whereas the middle and lower images show ovaries and spermathecae from corresponding female counterparts. Black lines visible in the spermathecae indicate the circular orientation of spermatozoa. Red boxes highlight regions of ovarian development. The deformed or broken appearance of the spermathecae is due to dissection artefacts rather than temperature effects. Scale bar = 100 µm.

To identify the reproductive organs affected by thermal stress and investigate the potential mechanism underlying the fertility effects, we dissected the reproductive organs of both treated and untreated adults (Figures 1B and 2B). Figure 1B showed that the ovaries of unfertilised females appeared hollow and lacked distinct structures, whereas fertilised ovaries displayed obvious rounded egg structures. Ovaries of females exposed to 38, 4 and -1°C appeared visually similar to those of fertilised females, with slight deformities. There is a bigger visibility difference when looking at the treated and untreated spermathecae, where the absence of black lines (spermatozoa; (Pascini et al., 2013) can be seen when the females are not fertilised and obvious black lines when they have been mated and fertilised. The spermathecae of treated females exhibited faint black lines or, in some cases, no visible black lines at all.

Although treated and untreated male accessory glands appeared similar morphologically (Figure 2B) only minor differences in size were observed not sufficient to confidently attribute an effect to thermal stress. To further investigate the sterility observed in treated males, the reproductive organs of their female counterparts were dissected and examined. Examination of the female counterparts mated to treated males revealed ovaries and spermathecae that more closely resembled those of unfertilised females, suggesting impaired fertilisation or sperm transfer.

### Distribution map

Figure 3A is based solely on the thermal limits (LT90) without incorporating other ecological niche constraints. Large regions at higher latitudes in North America, Europe and some parts of northern Asia were classified as too cold for survival, while small parts of Europe and southern parts of South America were identified as too cold for reproduction. In contrast, tropical and subtropical regions across South America, sub-Saharan Africa, Southeast Asia and northern Australia were classified as suitable for both survival and reproduction under the LT90 thresholds. Notably, regions in North Africa (including much of the Sahara Desert), and parts of South Asia were classified as too hot for reproduction and survival. The gap between survival and reproduction limits is clearly visible on the map in regions where temperature allows survival but not reproduction.

**Figure 3:**
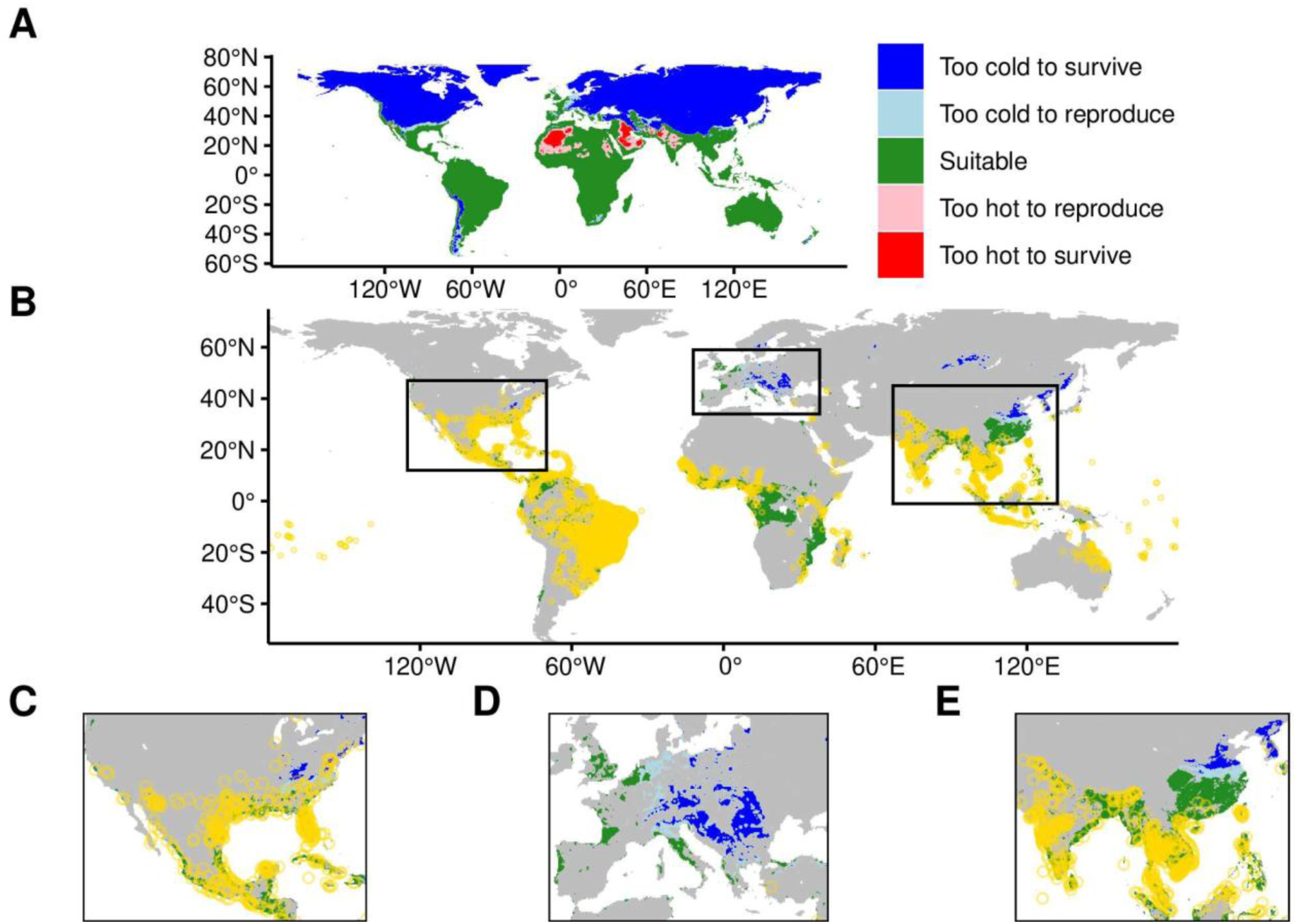
A) Predicted suitability of *Ae. aegypti* distribution based solely on experimentally derived thermal survival and fertility limits. Potential current habitat range of *Ae. aegypti* worldwide (B) and for North America (C), Europe (D) and East Asia (E) separately. Yellow circles indicate recorded occurrence locations of *Ae. aegypti*, overlaid for comparison with the experimentally predicted suitability map. Coloured regions represent suitability categories derived from LT90 cut-offs from the experimental data with areas labelled as too cold to survive (blue), to reproduce (light blue), suitable (green), too hot to reproduce (light red), and to survive (red).

Figure 3B-3E shows the current ecological niche of *Ae. aegypti*. The tropics are generally suitable which matches the distribution of *Ae. aegypti*, especially in South America, Africa, and Southeast Asia. Agreement between the predicted thermal niche and occurrence records was particularly strong across much of the Americas and Southeast Asia, although some regional discrepancies were evident. Suitable conditions were predicted across large areas of sub-Saharan Africa where relatively few occurrence records were present, whereas occurrence records in parts of the Indian subcontinent and the Black Sea region, including southern Russia and Georgia, extend beyond areas predicted to be suitable. Large parts of Central Europe appear generally suitable for the mosquito but are currently too cold in winter to survive or reproduce. Larger parts of China appear suitable too, but the mosquito has not spread northwards so far.

Figure 4 shows the ecological niche of *Ae. aegypti* with warming climates in the 2050s. While the tropics are still suitable for the mosquito, some parts of Central Europe and temperate US that have previously been too cold to reproduce in winter will become suitable too. Parts of India on the other hand might become too hot in summer for successful reproduction by the mosquito.

**Figure 4:**
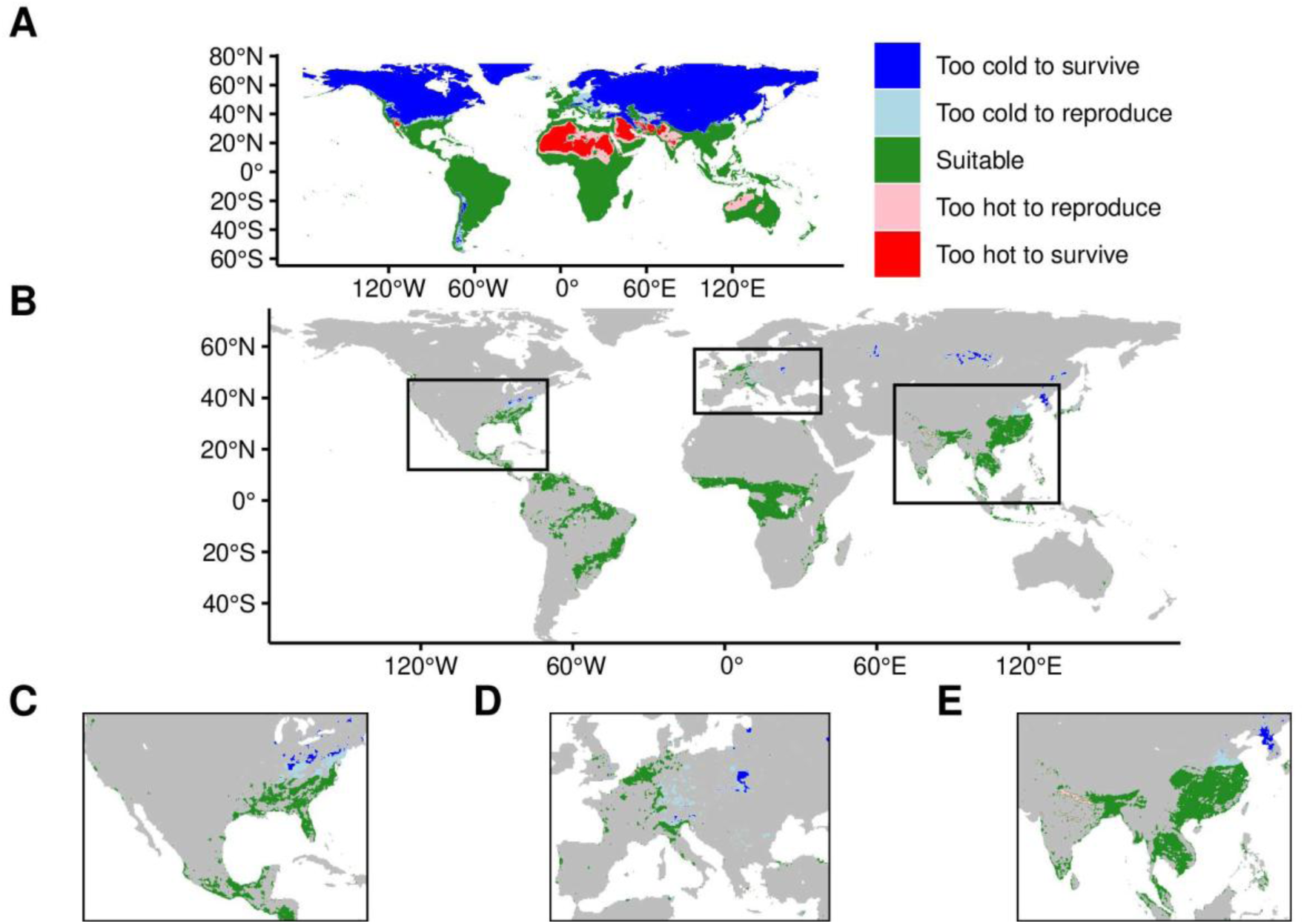
Ecological niche of *Ae. aegypti* with warming climates in the 2050s. Potential future habitat range of *Ae. aegypti* worldwide (A) and for North America (B), Europe (C) and East Asia (D) separately.

## DISCUSSION

Temperature extremes are well known to affect insect physiology and fitness. In this study, we found that *Aedes aegypti* has a significantly more restrictive thermal fertility limit (TFL) compared to its critical thermal limit (CTL), meaning mosquitoes may survive temperature extremes but lose the ability to reproduce. These findings highlight the importance of considering sublethal reproductive constraints when assessing thermal tolerance and species fitness under varying temperature regimes. We also observed significant morphological differences in female reproductive organs following thermal stress, providing insight into the mechanisms underlying fertility loss.

Our findings are consistent with Parratt et al. (2021), who observed fertility loss in *Drosophila* species at temperatures below upper lethal thresholds, with a maximum heat side gap of 4.3°C between TFL and CTL. At the upper thermal margin in *Ae. aegypti*, we observed fertility loss occurring at 9°C (males) and 5°C (females) below the corresponding CTL, supporting the concept introduced by Walsh et al. (2019) that fertility limits offer a more sensitive measure of thermal stress than survival-based limits alone. Previous studies similarly show that *Ae. aegypti* can remain active and survive at high temperatures while experiencing substantial reductions in fecundity (Rowley and Graham, 1968; Villareal et al., 2017; Brady et al., 2013; Ziegler et al., 2022; Delatte et al., 2009, Goindin et al., 2015). Notably, temperatures exceeding the observed TFL of 38°C have already been recorded during recent European heatwaves (The Copernicus Climate Change Service, 2024), suggesting that reproductive failure may occur under naturally occurring environmental conditions even when adult survival is maintained. Thus, at the upper thermal margin, incorporating TFL may further restrict predicted suitability in already warm regions. However, the implications of warming at the lower thermal margin may be particularly relevant for Europe. Under projected warming for the 2050s (SSP2-4.5), parts of Central Europe that are currently too cold for winter reproduction are predicted to become suitable, potentially increasing opportunities for establishment as temperature cross the lower thermal fertility threshold.

In addition to high temperatures, we also observed fertility loss following exposure to low temperatures. Approximately 80% of male and female adults survived a 6-hour exposure at 4°C, consistent with previous studies demonstrating adult survival following short-term cold stress (Culbert et al., 2019; Kramer et al., 2021). However, fertility declined at substantially higher temperatures than those causing mortality, indicating that reproductive performance is more sensitive to cold stress than survival. This is consistent with Culbert et al. (2018), who reported declining fertility at low temperatures in a Brazilian colony. Together, these findings suggest that thermal fertility limits provide a more sensitive measure of cold stress than survival alone.

To investigate the mechanisms underlying reduced fertility, we examined the reproductive organs of both males and females. Females exposed to extreme temperatures exhibited distinct morphological changes in the ovaries and spermathecae, while females mated to treated males displayed reproductive organ morphologies resembling those of unfertilised females. These findings suggest that thermal stress primarily affects process associated with successful insemination and sperm storage, as evidenced by abnormalities in the spermathecae, rather than causing substantial disruption to ovarian morphology. or fertilisation. These findings are consistent with those of Bader and Williams (2012), who reported similar reductions in reproductive performance following thermal stress associated with impaired ovarian development and egg production in mosquitoes. Although no obvious morphological differences were observed in male accessory glands, this does not exclude the possibility of functional impairment. Thermal stress may affect sperm production, sperm viability, sperm transfer, or seminal fluid composition without causing visible structural changes (Wolfner et al., 2007; Scolari et al., 2012; Sirot et al., 2011). Male accessory glands produce seminal fluid proteins (SFPs) that regulate post-mating responses in female mosquitoes, including blood-feeding behaviour and oviposition (Catteruccia et al., 2009; Gillott, 2003; Avila et al., 2011). Therefore, disruption of seminal fluid protein function remains a possible explanation for the reduced fertility observed in this study. However, the present data do not allow us to distinguish between effects on seminal fluid proteins, sperm function, or other aspects of reproduction.

Figure 1 shows a clear divergence between survival (CTL) and fertility limits (TFL), particularly at lower temperatures, where adults may survive short exposures but remain reproductively impaired. When integrated into niche models, this suggests that some regions may intermittently support survival and, during favourable periods, limited reproduction, without necessarily supporting long-term population persistence. This interpretation is consistent with observations that *Ae. aegypti* has appeared at the margins of its historical range, including southern Europe (Kraemer et al., 2015; ECDC, 2023), and that climatic suitability is expected to shift under warming scenarios (Ryan et al., 2019; Liu-Helmersson et al., 2019). In temperate regions, seasonal asymmetry is likely critical, with summer conditions permitting reproduction even where winter temperatures constrain overwintering. Models based solely on survival thresholds may therefore oversimplify these dynamics by classifying such areas as uniformly unsuitable, despite periods when reproduction remains possible. Incorporating fertility responses helps distinguish regions that support temporary survival and reproduction from those capable of sustaining long-term establishment. For example, although some regions such as the United Kingdom may periodically support adult survival and reproduction, this should not be interpreted as evidence of long-term population persistence.

“All models are wrong, but some of them are useful” – George Box, 1976. One of the key limitations of our model is its inability to fully capture fine-scale environmental variation and temporal exposure dynamics. Microclimates can result in substantial deviations from ambient air temperatures, particularly in sheltered or indoor environments where *Aedes aegypti* is commonly found (Rodrigues et al., 2015; Lembrechts et al., 2018). In addition, our model is based on monthly temperature extremes and does not account for short-duration thermal events, such as heatwaves or cold snaps, that may disproportionately affect fertility relative to survival. Given that our experimental exposures are conducted over defined short periods, these transient conditions may be particularly important in determining real-world reproductive outcomes. Incorporating higher temporal and spatial resolution environmental data will therefore be important for refining predictions of where and when populations may persist. Additional limitations of this study include focus on a single life stage and the use of a laboratory strain. It is well-established that other life stages are also sensitive to extreme temperatures, which can induce sterility. For instance, exposure to both low and high temperatures during the larval and pupal stages have been shown to cause sterility in adult *Drosophila* (Walsh et al., 2021; Vollmer et al., 2004). Future studies should determine TFL50 values across all life stages to provide a more comprehensive estimate of fertility loss throughout the mosquito life cycle. Although the observed responses broadly align with published data from field populations, laboratory-adapted mosquitoes may exhibit altered thermal tolerance. Future studies should therefore test field strains across similar temperature regimes. Importantly, thermal fertility limits alone do not predict disease risk. Vector competence and pathogen development also respond to temperature. Incorporating both mosquito and pathogen thermal biology is critical for future risk models.

Our findings demonstrate that incorporating thermal fertility limits (TFL) moderately improves the predictive accuracy of temperature performance models beyond those based solely on critical thermal limits (CTL). These findings emphasize that reproductive capacity, rather than survival alone, more accurately defines the thermal tolerance boundaries and fitness potential of species under variable temperature regimes.

