## Supplementary Information for "Thermal fertility limits reveal underestimated climatic suitability for *Aedes aegypti* in temperate regions"

Methodology

**Gaussian equation**

$$Proportion\ of\ survival\ or\ sterility = a \times exp\left(-\left(\frac{(Temperature - m)}{s}\right)^{b}\right)$$

a = Maximum predicted proportion of survival or sterility

m = Temperature at which survival or sterility is maximal

s = Sharpness of curve

b= Steepness of curve

**Map**

We use the biomod2 package in R. We check the three models RF (random forest), GBM (generalised boosted regression), and MAXENT and use a 70:30 split for training and testing data to avoid overfitting. RF shows the best fit (mean PR AUC 0.88, GBM: 0.83, Maxent 0.8) but can still lead to overfitting in some cases. Based on response curves for predictor variables, we use the mean ensemble prediction of all three models.

According to Barbet-Massin et al. (2012), we generate the same number of pseudo-absences as we have presence data points. The used resolution results in 1080 times 2160 grid cells globally, of which about 30% are land covered. The 3,969 occurrence point thus cover approximately 0.6% of the land grid cells, and thus 10 repetitions of PA sampling should be suitable. However, we observe that this high number of runs again leads to overfitting and here we only use 3 repetitions.

We use *Ae. aegypti* occurrence data from Kraemer et al (2015). The data set contains 19,929 data points. To avoid oversampling, we filter the data set by only retaining the first data point within a 25 km radius, resulting in 3,969 occurrence points. We use gridded climatic and elevation data with a resolution of 0.3° (roughly 18 km) from WorldClim v2.1. Monthly Relative Humidity (RH) is calculated from monthly average temperature (T) and water pressure (Vp) using the Arden Buck equations for temperatures below and above freezing. We use minimum values for RH and precipitation as these show less multicollinearity with other variables than mean values. Human population data is available for 2020 from GPWv4-UN-adjusted and is regridded to 0.3°. Monthly Normalized Difference Vegetation Index (NDVI) data for 2022 is available from AVHRR GIMMS-3G+. The annual average is taken and regridded to 0.3°. We do not use temperature as a predictor because we then constrain the found ecological niche with the temperature data from the lab. We did check using annual temperature as an additional or sole predictor, see below plots.

We use the calculated LT90 cut-offs to identify areas that are too hot and too cold to live or reproduce. Using WorldClim v2.1 data sets for maximum (bio_wc5) and minimum (bio_wc6) temperatures, we mask unsuitable areas. The resulting map shows areas as too cold to survive (blue), to reproduce (light blue), suitable (white), too hot to reproduce (light red), and to survive (red), for males and females. Constraining the found ecological niche with lab data increases the predictive performance of the model, measured as F1 score from 0.76 to 0.77.

For future climates, we use bias-corrected climate projections for the SSP2-4.5 scenarios from the NEX-GDDP-CMIP6 data set, including humidity, precipitation, and wind. We use chose three models: ACCESS-CM2 (Australia), NorESM2-MM (Norway), MPI-ESM1-2-LR (Germany) to capture a range of climate sensitivities. The Monthly averages are calculated from daily data for the period 2050 to 2059 to align with the WorldClim data format. Similarly, bio_wc5 and bio_wc6 are calculated with climate projections for minimum and maximum temperatures to get LT90 cut-offs for the 2050s.

Supplementary Figure 1.1: Survival and sterility against high and low temperature. A) Males survival and hatching rate calculated using the (number of eggs/total number eggs). B) Females survival and hatching rate calculated using the (number of eggs/total).

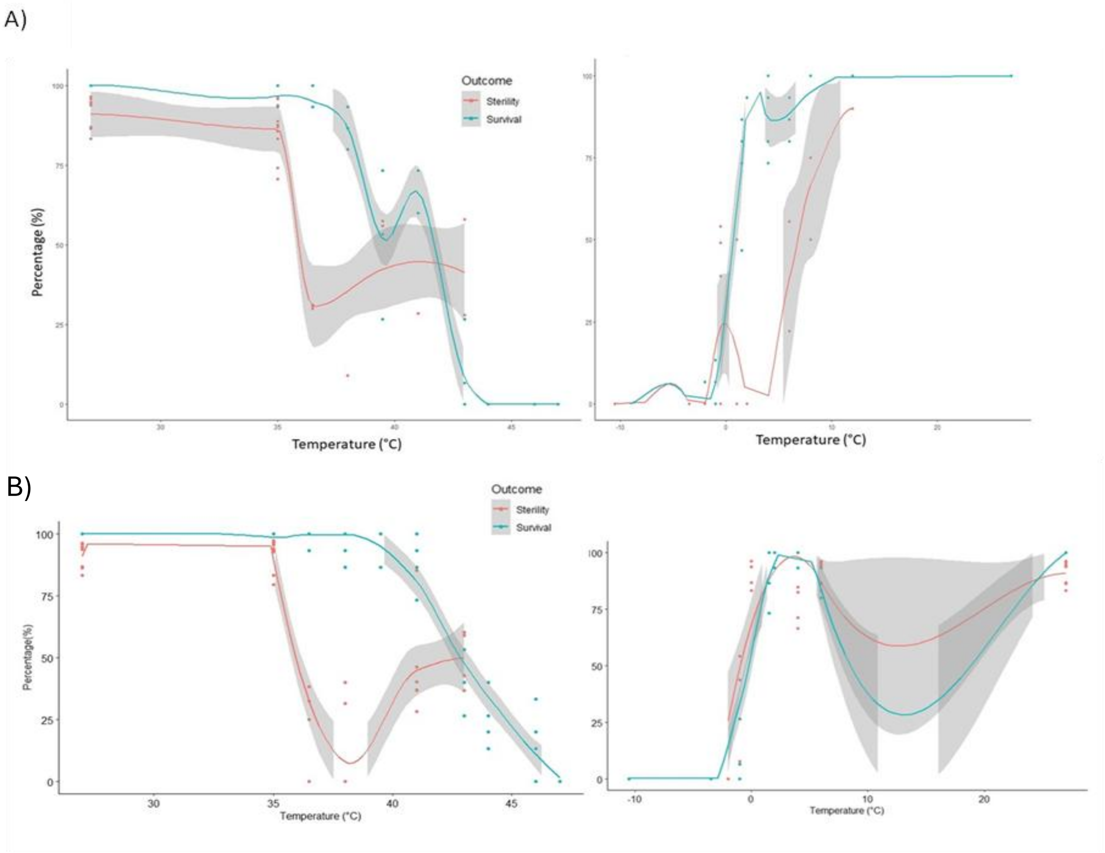

Supplementary Figure 2.1:

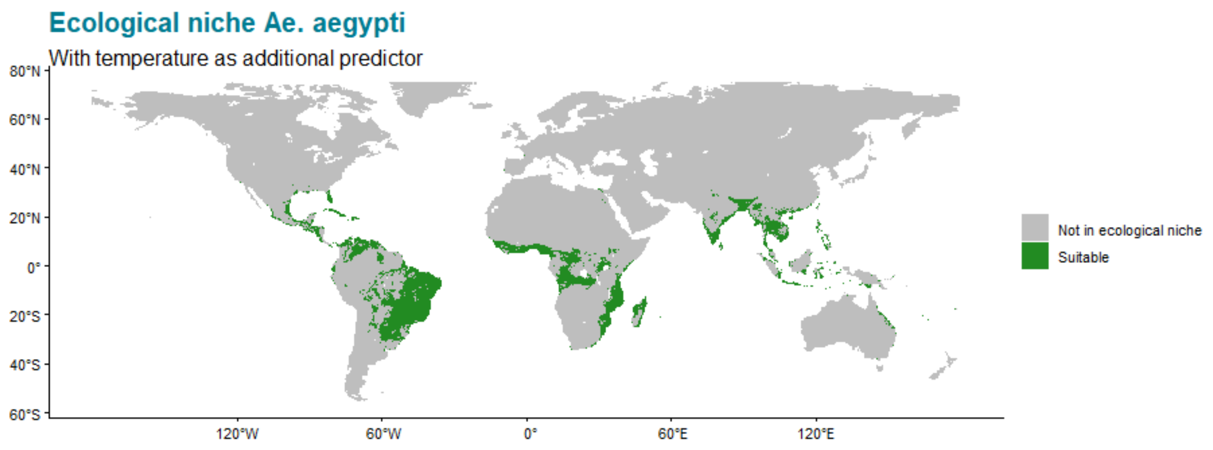


Supplementary Figure 2.2:

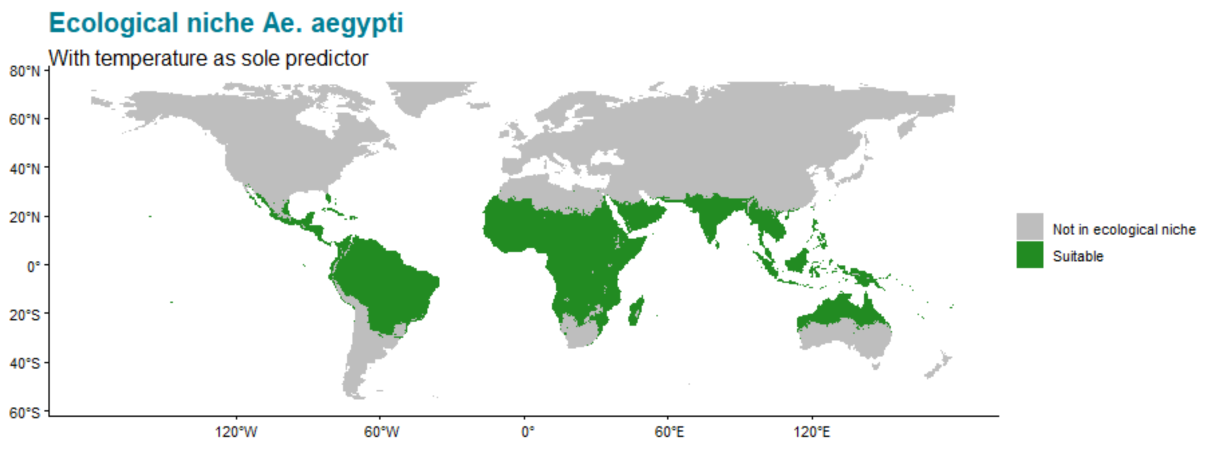


Supplementary Table 1

| Category | Proportion | Count |
|---|---|---|
| Too cold to survive | 0.4% | 14 |
| Too cold to reproduce | 0.3% | 10 |
| Not in ecological niche | 18.5% | 718 |
| Suitable | 80.6% | 3123 |
| Too hot to reproduce | 0.3% | 11 |
| Too hot to survive | 0% | 0 |
